# Development and evaluation of a wearable peripheral vascular compensation sensor in a swine model of hemorrhage

**DOI:** 10.1101/2023.02.20.529156

**Authors:** Francesca Bonetta-Misteli, Toi Collins, Todd Pavek, Madison Carlgren, Antonina Frolova, Leonid Shmuylovich, Christine M. O’Brien

## Abstract

Postpartum hemorrhage (PPH) is both the leading and most preventable cause of maternal mortality. PPH is currently diagnosed through visual estimation of blood loss or vital sign analysis of shock index (ratio of heart rate to systolic blood pressure). Visual assessment underestimates blood loss, particularly in the setting of internal bleeding, and compensatory mechanisms stabilize hemodynamics until hemorrhage is massive, beyond the point of pharmaceutical intervention. Quantitative monitoring of hemorrhage-induced compensatory processes, such as the constriction of peripheral vessels to shunt blood to the central organs, may provide an early alert for PPH. To this end, we developed a low-cost, wearable optical device that continuously monitors peripheral perfusion via laser speckle flow index (LSFI) to detect hemorrhage-induced peripheral vasoconstriction. The device was first tested using flow phantoms across a range of physiologically relevant flow rates and demonstrated a linear response. Subsequent testing occurred in swine hemorrhage studies (n=6) by placing the device on the posterior side of the swine’s front hock and withdrawing blood from the femoral vein at a constant rate. Resuscitation with intravenous crystalloids followed the induced hemorrhage. The mean LSFI vs. percent estimated blood volume loss had an average correlation coefficient of −0.95 during the hemorrhage phase and 0.79 during resuscitation, both of which were superior to the performance of the shock index. With continued development, this noninvasive, low-cost, and reusable device has global potential to provide an early alert of PPH when low-cost and accessible management strategies are most effective, helping to reduce maternal morbidity and mortality from this largely preventable problem.

## Introduction

Postpartum hemorrhage (PPH) is the leading cause of maternal mortality globally, accounting for 27% of maternal deaths worldwide.^1^ In the United States, PPH occurs in over 100,000 deliveries annually^2^ and is responsible for 11% of maternal deaths. Alarmingly, a national study revealed that black women have a 2.5 times higher pregnancy-related mortality ratio caused by PPH compared to white women.^3^ Beyond maternal death, complications caused by hemorrhage are responsible for the majority of *severe* maternal morbidities.^4–6^ Fortunately, early diagnosis and treatment significantly reduce mortality and morbidity associated with PPH. ^7–9 10–12^

The principal cause of PPH is uterine atony, in which the uterus fails to contract and tamponade vessels that previously perfused the placenta at a rate of 600-700 mL blood/min.^13, 14^ Treatment options in early stages of hemorrhage include pharmacologic agents, intrauterine-balloon tamponade, and low-level intrauterine vacuum to aid uterine contraction^15^. Later stages of hemorrhage require blood transfusion and surgical interventions.^16^ In most low-resource settings and in small US hospitals, blood stores and capacity for surgical intervention are limited.^17^ As a result, interventions for PPH are confined primarily to relatively cheap and easily accessible pharmacologic agents. However, these treatments are most effective at the onset of PPH – underscoring the importance of early diagnosis and treatment.

While early detection of PPH is critical, it is also challenging. The most common method for PPH diagnosis is visual estimation of blood loss due to its simplicity and zero added cost. However, it has been shown to underestimate blood loss up to 75%, with the amount of error increasing with increased PPH severity.^18, 19^ In addition to visual estimation of blood loss, providers may rely on vital signs such as heart rate, respiratory rate, and blood pressure. However, in early hemorrhage, compensatory mechanisms are activated to ensure adequate perfusion of vital organs, and in turn, vital signs remain stable. A primary example is peripheral vasoconstriction, which effectively shunts blood flow from the limbs to vital organs.^20, 21^ Several tools have been developed to help quantify blood loss^22, 23^ and estimate the degree of hemorrhage.^24, 25^ However, most are either limited in their ability to detect occult blood loss, which is common in uterine atony, or are only effective in settings of massive PPH (>1500 mL blood loss) when blood replacement products and surgical intervention are required. As such, there is an unmet need for a continuous monitoring method for PPH that is inexpensive, accessible, and sensitive to both external and internal bleeding.

We hypothesized that a light-based perfusion measurement, the laser speckle flow index (LSFI), could capture the peripheral vasoconstriction compensatory response, and thereby detect early indicators of hemorrhage. LSFI uses a laser and a camera to detect and quantify spatial and temporal changes in the speckle patterns to extract flow information. LSFI is linearly proportional to blood flow,^26–28^ and wearable laser speckle imaging sensors have been developed to provide real-time monitoring of perfusion and cardiovascular physiology.^29, 30^ Furthermore, LSFI has a high signal-to-noise ratio (SNR) and is minimally affected by skin color,^31^ providing excellent signal quality.^29^ We have developed a custom wrist-worn wearable LSFI sensor to detect early hemorrhage-associated changes in peripheral perfusion. The device is small, simple, battery-powered, and transmits data wirelessly via Bluetooth to track results from a mobile device such as a tablet. Such characteristics are ideal for patient monitoring, as evidenced by the pulse oximeter, a light-based device used globally for patient monitoring. We have characterized the sensor’s optical performance, validated a linear response from flow phantoms across a range of flow rates, and demonstrated utility as an early detection tool for hemorrhage in a swine model of controlled volume hemorrhage.

## Results

### 1. Custom wearable LSFI sensor design and operation

A small LSFI system was developed with a wearable “wrist-watch” form that continuously and non-invasively monitors peripheral perfusion for early detection of hemorrhage. Software controls the laser module and camera sensor, performs real-time video capture and data processing, and transfers data via Bluetooth to external devices (Figure 1).

**Figure 1.**
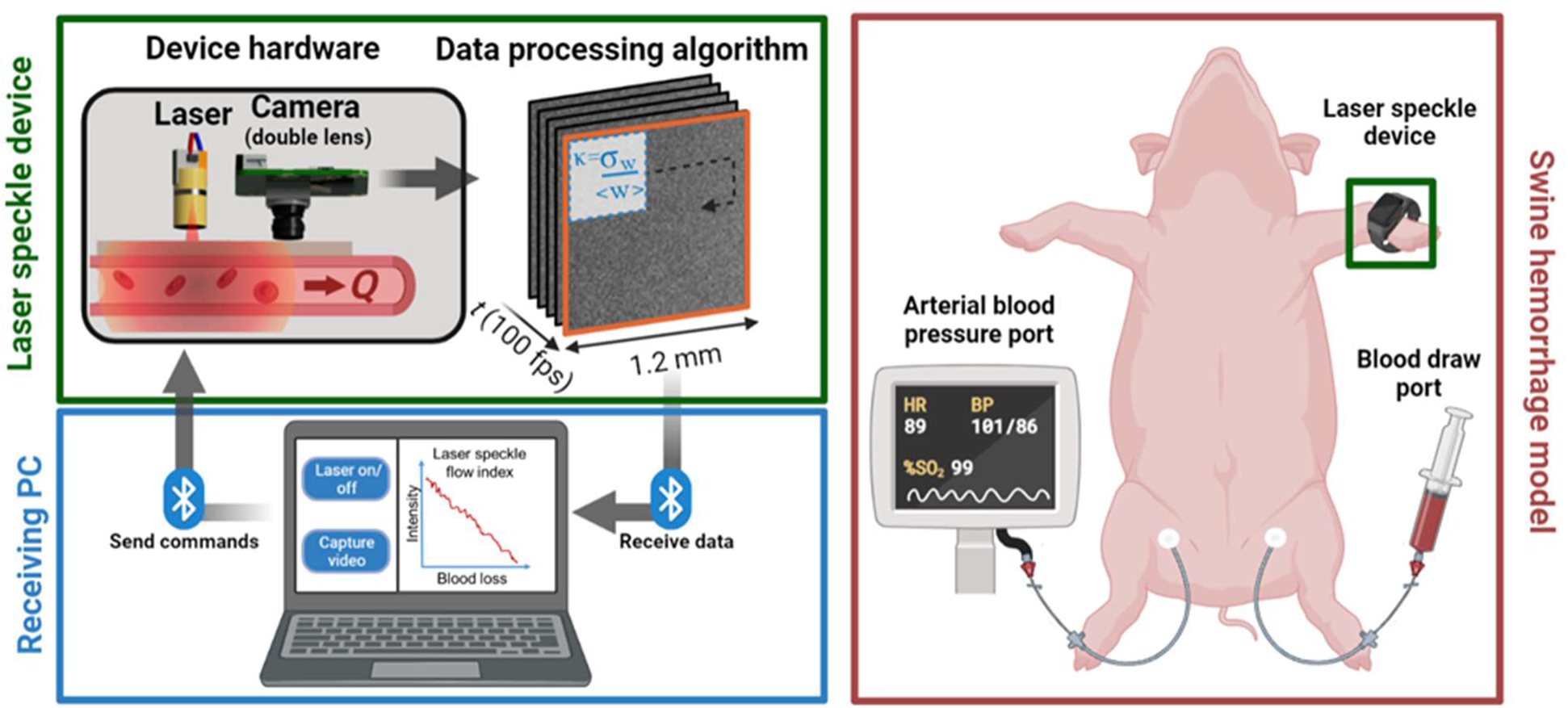
Data acquisition and visualization via Bluetooth communication used in swine studies. Left) User inputs to control laser and video capture are sent to the Raspberry Pi. Images are recorded and processed on the device and sent back to the computer where they are automatically plotted. Right) Experimental setup for swine hemorrhage model, with blood draw port and arterial blood pressure port. Device was worn on the swine’s “wrist”.

### 2. Impact of camera system parameters on laser speckle

The custom double lens Raspberry Pi camera system characteristics for three recording modes: 30, 80, and 100 frame per second (fps), are outlined in Table 1. The focal length of the single lens system is 3.04 mm and the f/# is 2.0. As the first inverted lens acts simply as a collimator, the focal length and f/# of the fixed focus lens can be used to describe the double lens system. The aperture size is calculated to be 1.52 mm, consistent with the physical measurement of the aperture.

**Table 1.** Calculated double lens system camera parameters.

| Method | 30 fps 1920p | 80 fps 640p | 100 fps 320p |
| --- | --- | --- | --- |
| Focal length | 3.04 mm | 3.04 mm | 3.04 mm |
| Lens diameter | 1.52 mm | 1.52 mm | 1.52 mm |
| f/# | 2.0 | 2.0 | 2.0 |
| Camera pixel size | 1.12 x 1.12 $\mu\text{m}$ | 2.24 x 2.24 $\mu\text{m}$ | 4.48 x 4.48 $\mu\text{m}$ |
| Distance per pixel | 1.12 $\mu\text{m}$ | 2.24 $\mu\text{m}$ | 4.47 $\mu\text{m}$ |
| Magnification | 1.000x | 1.000x | 1.002x |
| Speckle size | 7.73 $\mu\text{m}$ | 7.73 $\mu\text{m}$ | 7.73 $\mu\text{m}$ |
| Speckle:Pixel Ratio | 6.90 | 3.45 | 1.72 |
| Spatial resolution | 2.19 $\mu\text{m}$ | 2.5 $\mu\text{m}$ | 3.9 $\mu\text{m}$ |

For images captured at the default 30 fps settings, the pixel size is 1.12 µm with no pixel binning, however, at the default 80 and 100 fps settings, the pixel size is 2.24 µm and 4.48 µm due to 2x and 4x binning, respectively. The object distance per pixel calculated from 30, 80, and 100 fps video captures was 4.47, 2.24, and 1.12 µm, respectively. As expected, this resulted in equivalent magnifications of ∼1.0x under each condition. The calculated speckle size was 7.73 µm (Eqn. 3 in Methods), which was equal for all modes. Thus, for the 30, 80, and 100 fps imaging rates a speckle-to-pixel ratio of 6.90, 3.45, and 1.72 was calculated, respectively. At 100 fps, the Nyquist criterion was not achieved, while the two lower sampling rates surpassed Nyquist. The three methods, from lowest to highest fps, produced different resolutions: 2.19, 2.5, and 3.9 µm, respectively.

### 3. Laser Characterization

The low-cost laser spectrum used in the wearable device was stable over a six-hour monitoring period and the performance of this laser was comparable to that of a highly coherent and stable benchtop laser that served as a benchmark (Figure 2a-b). The benchmark laser had a peak wavelength of 785.12 nm at room temperature and varied within a range of 0.607 nm. The peak wavelength of the low-cost laser was 791.25 nm at room temperature, and the variation was within a range of 0.605 nm if temperature was held constant (Supplemental Figure 1).

**Figure 2.**
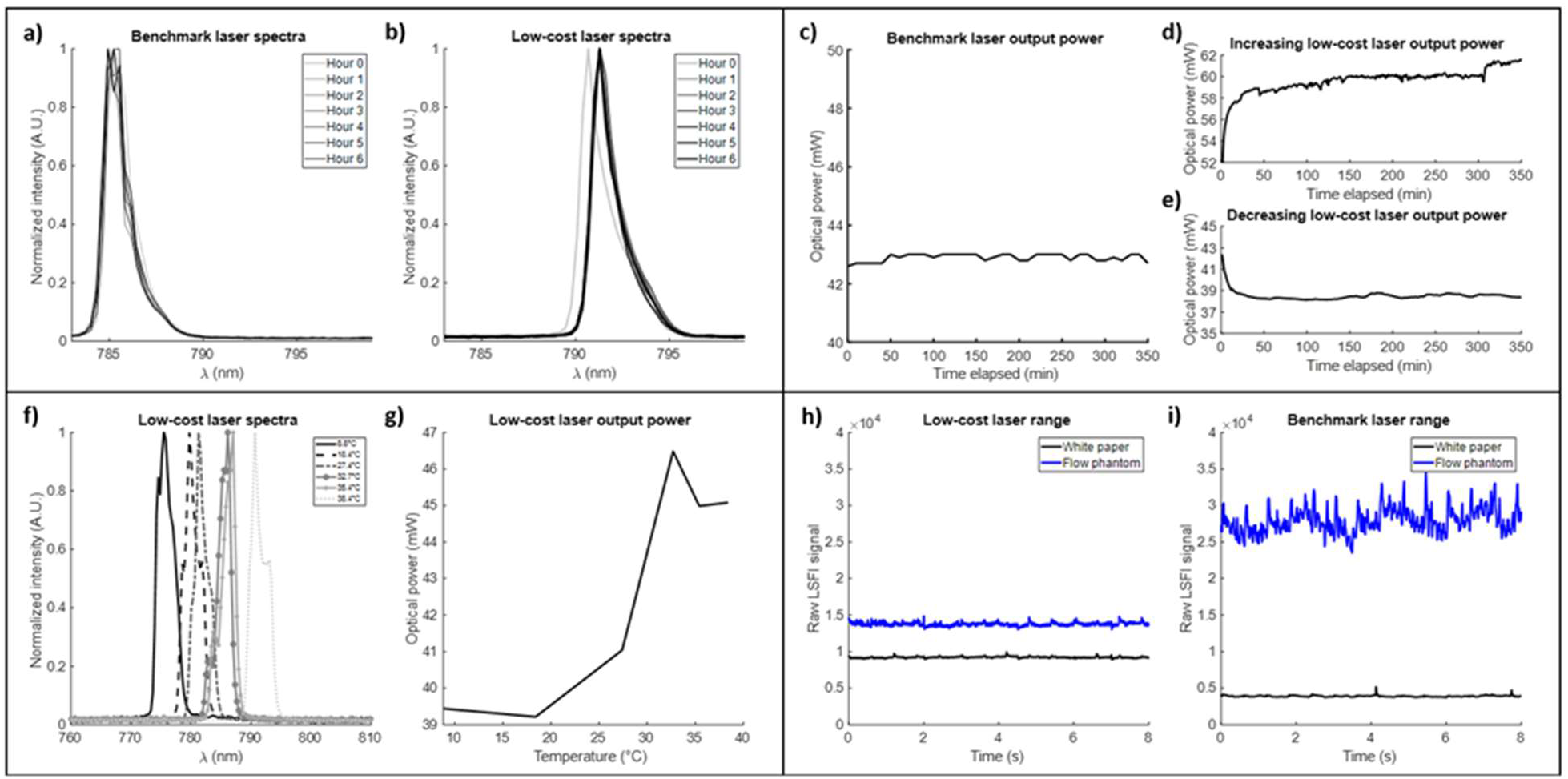
Laser characterization. Overlayed spectra of the benchmark (a) and low-cost laser (b) every hour for six hours. Optical power output of the benchmark laser (c), an initially increasing low-cost laser (d), and an initially decreasing low-cost laser (e). Low-cost laser spectra (f) and optical power output (g) over temperatures ranging from 8.8-38.4°C. Range of LSFI values produced by benchmark laser (h) and low-cost laser (i) while measuring static and moving media at 5 ms exposure time.

During a six-hour period, the output optical power of the benchmark laser and multiple low-cost lasers were recorded. The benchmark laser produced an average output power of 42.90 mW and varied within a range of 0.40 mW or 0.9% (Figure 2c). A batch of identical low-cost lasers from a single vendor were tested, and some decreased in the initial twenty-minute period, while others increased (Figure 2d-e). The optical power of the low-cost laser had a percent change of 13.8% and 8.6% for the increasing and decreasing lasers, respectively, within the first twenty minutes of measurement. The lasers remained stable for the remaining period with a percent change of 6.3% and 0.9%, respectively. For the low-cost lasers, intra-laser variability was low, but inter-laser variability was high. Unique laser identifiers tracked the use of each of the characterized lasers for the subsequent experiments.

The low-cost laser does not come equipped with a temperature regulating driver, thus characterizing the impact of temperature on laser stability is crucial. Decreasing the temperature of the laser caused lower optical power output and a lower peak wavelength (Figure 2f-g).

The maximum and minimum LSFI values for each laser were characterized (Figure 2h-i) by measuring white paper and a flow phantom, respectively. Using the benchmark laser, the minimum and maximum LSFI values were 3,913 and 27,960, respectively, with a total range of 24,050. The low-cost laser system produced a minimum and maximum LSFI value of 9,241 and 13,820, respectively, with a range of 4,574. The benchmark laser produced a broader range of LSFI values and higher and lower absolute values in the static and dynamic conditions.

### 4. Device power consumption

The device consumed 0.85 W (170 mAh) during idle status and 2 W during full operation (410 mAh) (Table 2). The PiSugar battery has a maximum capacity of 1200 mAh, and thus the battery life will last approximately 3 hours when actively collecting data.

**Table 2.**
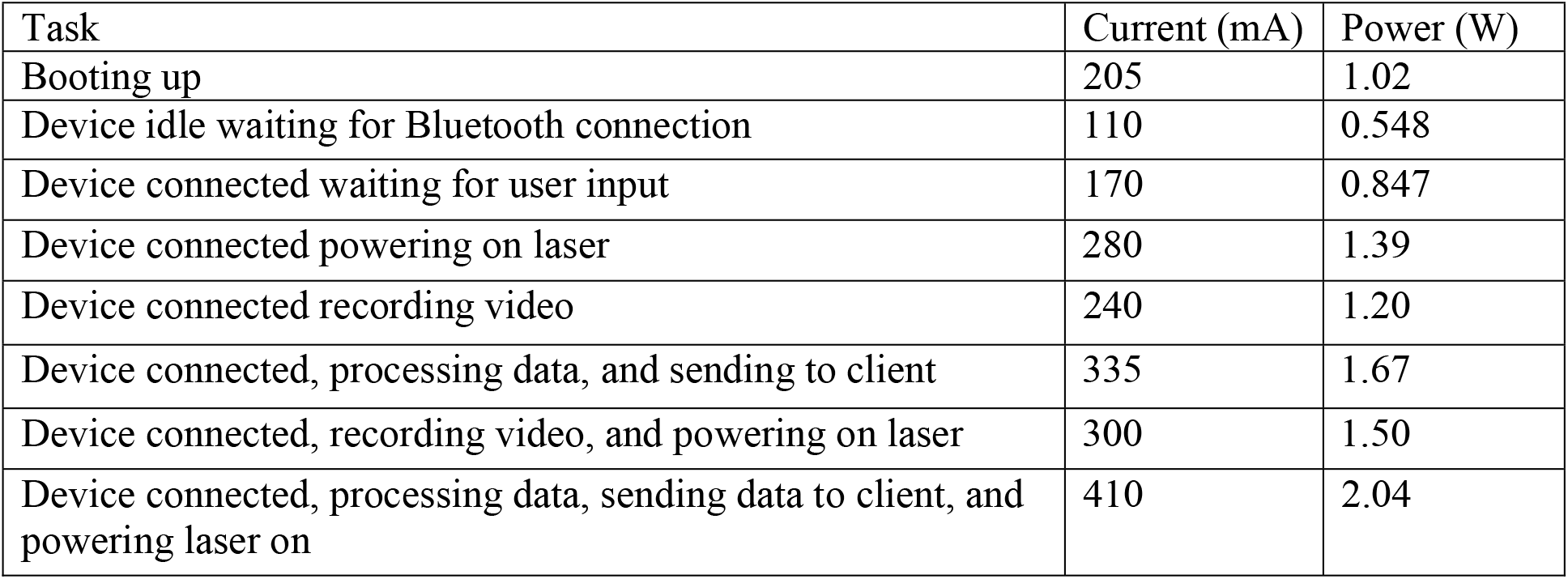
Power consumption of device performing various tasks related to data capture and processing; voltage held constant at 4.98 V.

### 5. Device performance in vitro

As expected, the relationship between mean LSFI and fluid velocity was linear in the low-cost and benchtop laser systems, with a strong correlation coefficient of 1 and 0.99 achieved, respectively (Figure 3). In addition, the benchtop laser exhibited a steeper slope (8535) compared to the low-cost laser (6561).

**Figure 3.**
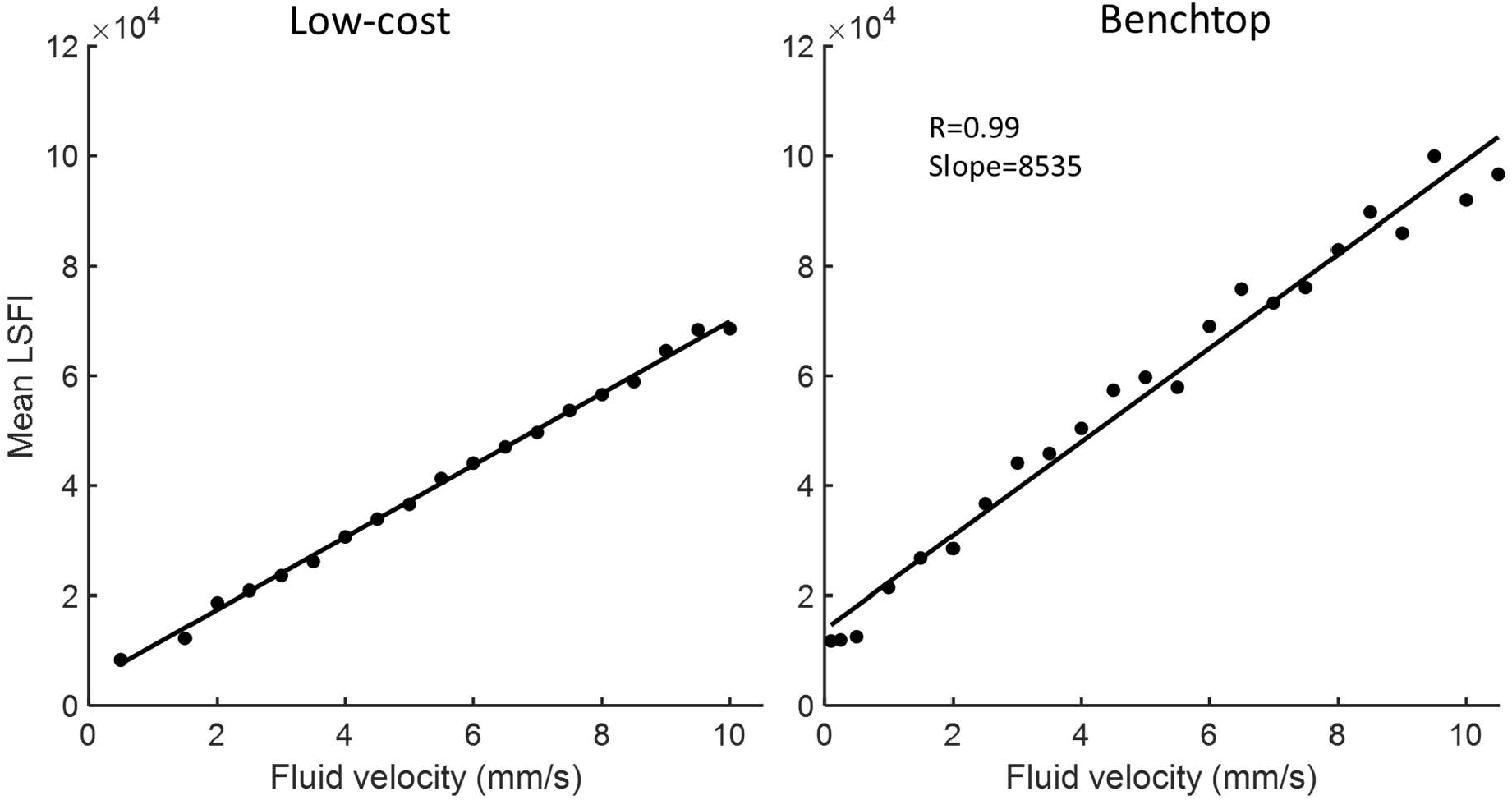
Mean LSFI measured while recording various physiological blood velocities with 0.5 ms exposure time.

### 6. Device performance in swine model of hemorrhage

Plots of the peak-normalized mean LSFI signal per minute of the study for each swine (Figure 4) displayed the relationship of each response to hemorrhage and crystalloid resuscitation. There was a distinct decrease in LSFI signal during periods of blood draw matching the rate of removal and an apparent increase in LSFI during subsequent crystalloid infusions in all models except swine 5. Unlike other swine, swine 5 was heparinized. Heparin has been reported to inhibit vasoconstriction, and therefore the swine’s ability to vasoconstrict in response to blood loss may have been affected.^32–35^ Due to this fact, swine 5 was not considered during subsequent quantitative analysis. At the start of blood draw, swine 1-6 had been given IV fluids volumes of 740, 870, 390, 564, 438, and 168 mL, respectively, which reflect differences in the length of time it took to prepare each swine for the experiment. Swine 1 experienced vein collapse in the left femoral vein towards the end of the blood draw procedure. The vein collapse was discovered as it became challenging to draw blood from the swine and was detected as a momentary increase in LSFI signal on the laser speckle device. Additionally, swine 4 experienced a blood clot during the crystalloid infusion phase, which was identified by the veterinary staff and was also detected by the laser speckle device as a sharp decrease in the LSFI signal.

**Figure 4.**
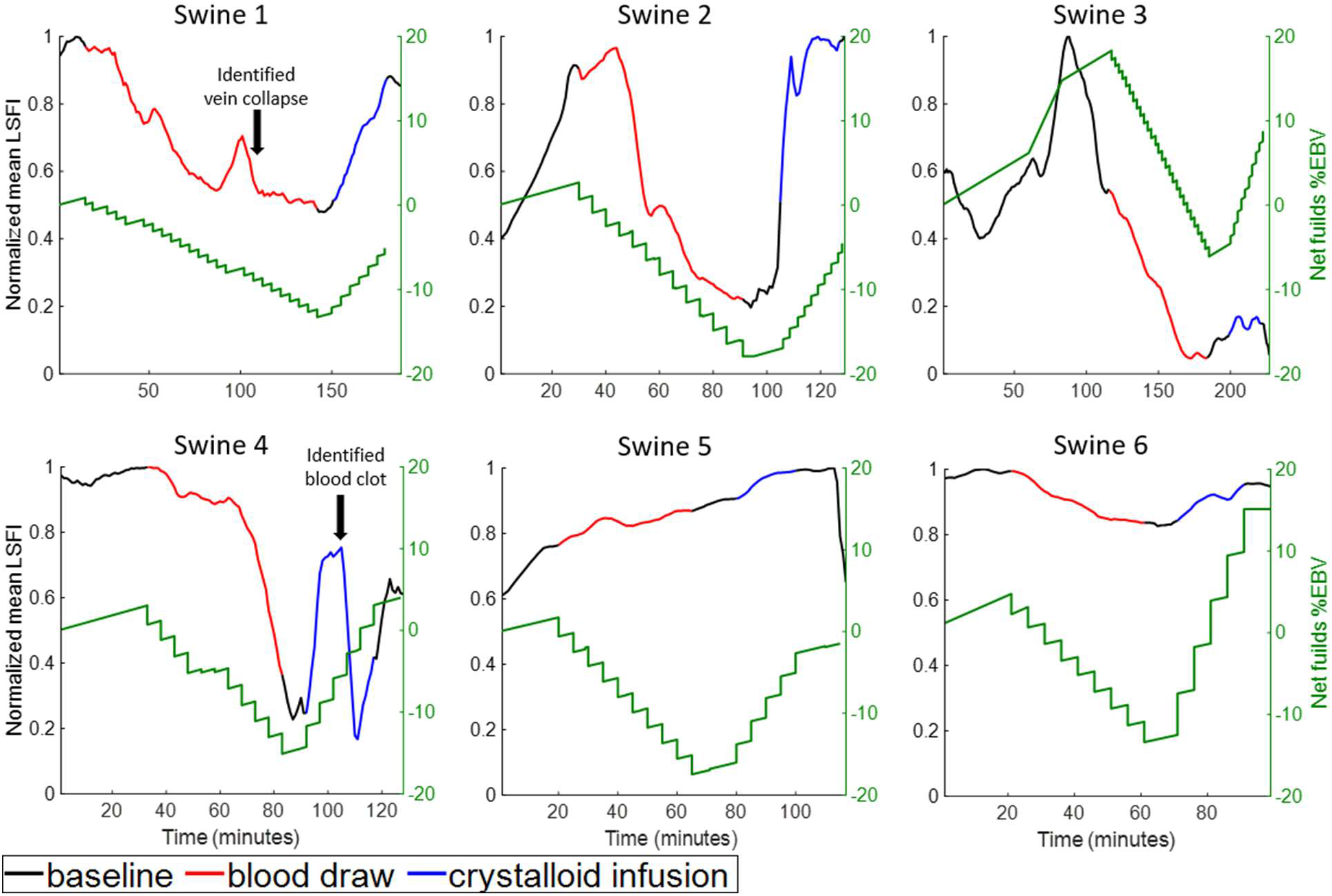
Peak-normalized mean LSFI signal during swine hemorrhage studies compared to net fluid change for each of six swine.

The baseline LSFI signal for swine 3 varied more than other swine during the alteration of isoflurane dosage (Supplemental Figure 2). This deviation may have been the result of the vasoactive properties of isoflurane as each swine may have a distinct response to the drug. Swine 3 received a much higher dose of isoflurane (3%) compared to other swine (1.5-2%). Other swine also required titration of the isoflurane dose throughout the procedure to prevent manual breathing.

Overlays of heart rate (HR), mean arterial pressure (MAP), and shock index (SI) with the peak-normalized mean LSFI signal over time displayed revealed minimal correlation between LSFI and HR, similar responses between LSFI and MAP, and an inverse response between LSFI and SI for each swine (Supplemental Figures 3-5). Swine were given oxygen throughout the procedure, resulting in highly stable blood oxygen saturation (SpO2) measures (Supplemental Figure 6). Continuous vital sign recordings were not available for swine 1; instead, measurements were recorded every 15 minutes. In addition, LSFI, MAP, HR, and SI were correlated with net fluid change during hemorrhage (Figure 5) and during resuscitation (Figure 6). LSFI was normalized by the initial signal at the start of hemorrhage and resuscitation, respectively. During hemorrhage, LSFI had the highest correlation of −0.94 with net fluid loss compared to other vital signs monitored. MAP had similar trends as LSFI with net fluid change of −0.92. HR had a correlation of 0.14, which was significantly worse than all other measures. SI had a correlation of 0.88 and an inverse trend as LSFI.

**Figure 5.**
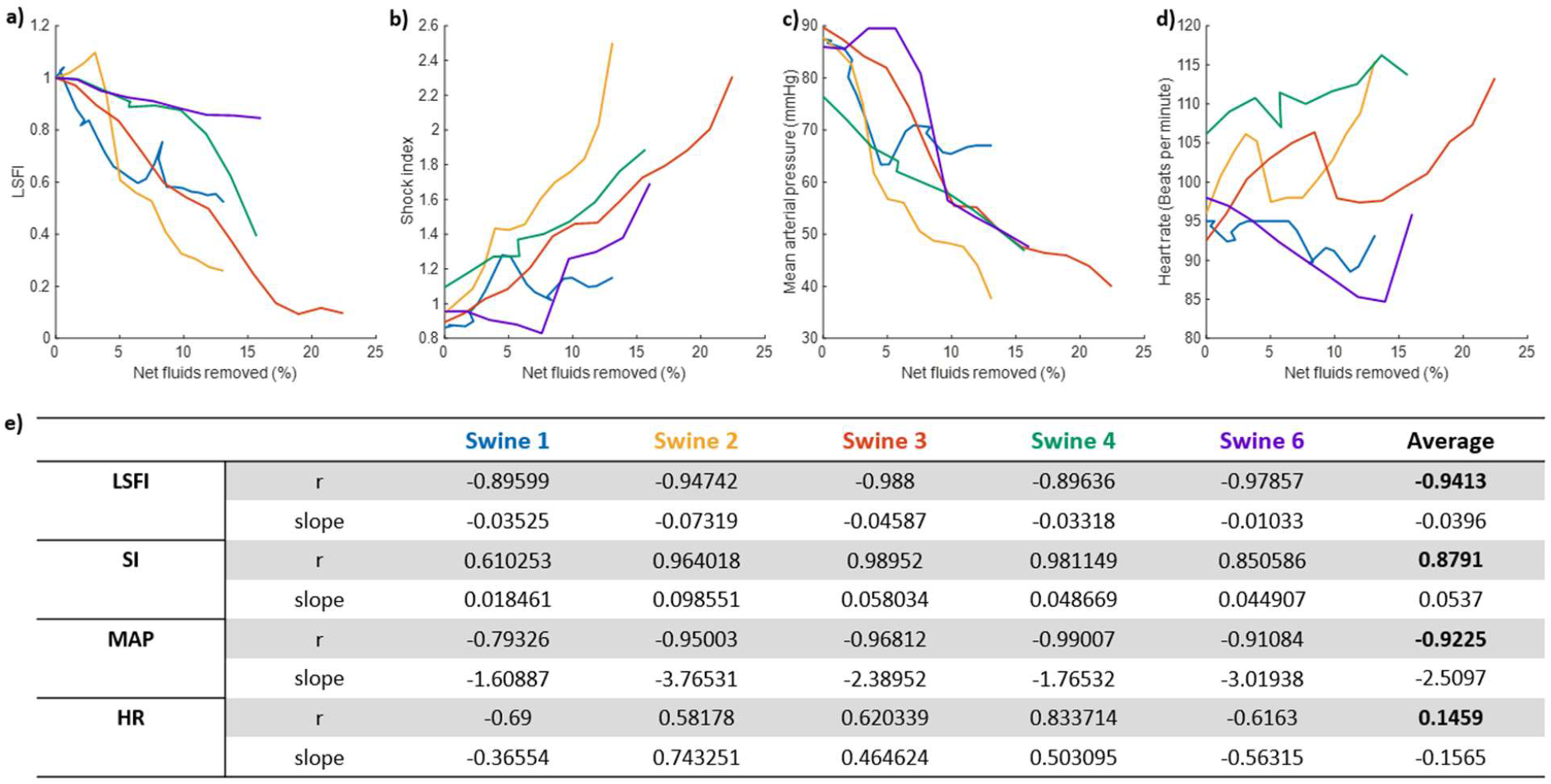
Response of LSFI sensor and vital signs during hemorrhage. Correlation between LSFI (a), shock index (SI) (b), mean arterial pressure (MAP) (c), and heart rate (HR) (d) vs net fluid change since the start of blood removal for each swine during blood loss only. (e) Summary of correlation coefficients and slopes for each case.

**Figure 6.**
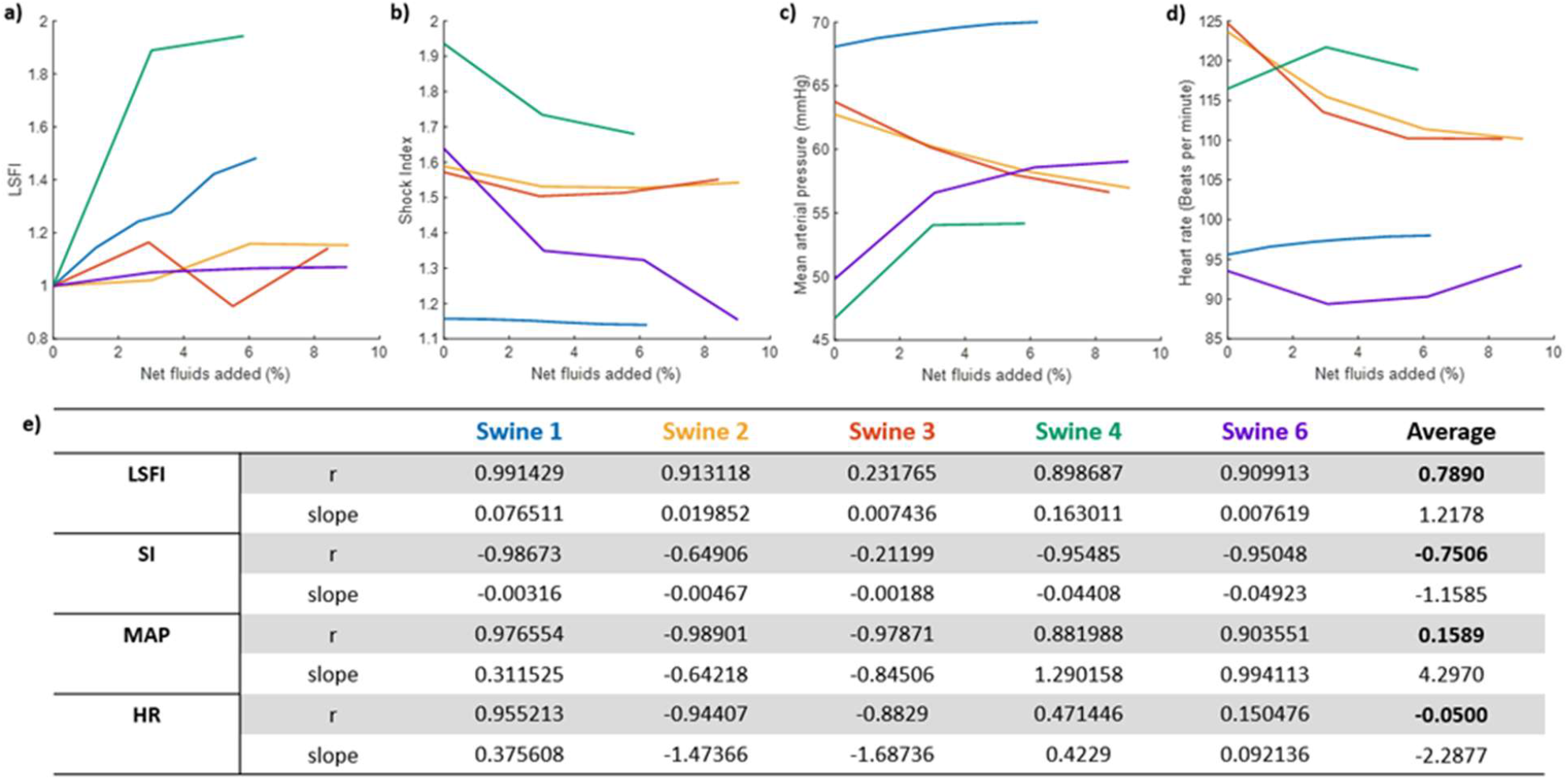
Response of LSFI sensor and vital signs during crystalloid infusion. Correlation between LSFI (a), shock index (SI)(b), mean arterial pressure (MAP) (c), and heart rate (HR) (d) vs. net fluid change in each swine since the start of crystalloid infusion. (e) Summary of correlation coefficients and slopes from linear fitting for each case.

During resuscitation, LSFI demonstrated a strong positive correlation with net fluid change (r=0.79), whereas MAP, HR, and SI had poorer correlation with net fluid change, with correlation coefficients of −0.75, 0.16, and −0.05 respectively (Figure 6). Data collected from swine 4 following the detected blood clot was omitted from this analysis.

We also tested the correlation between LSFI, MAP, and SI (Supplemental Figure 7). LSFI vs. MAP showed strong correlation during hemorrhage (r=0.93), but weak correlation during resuscitation (r=0.36). Similar findings were observed when performing a correlation of LSFI vs. SI during hemorrhage (r=-0.90) and resuscitation (r=-0.79).

### 7. Laser speckle video parameter analysis

The raw, unsmoothed LSFI signal produced throughout the hemorrhage procedure was down-sampled by frame rate from 100 fps to 50 and 10 fps. In addition, the frame size was varied from a full frame to one half- and one quarter-frames. The down-sampled frame rate and frame size of the entire swine hemorrhage study resulted in minimal changes in the mean LSFI trend result (Supplemental Figure 8).

## Discussion

Hemorrhage can be a devastating complication of not only childbirth, but also trauma, surgery, and bleeding disorders. Tools that can provide early indication of concerning blood loss could transform management from reactive to preventive, particularly in the postpartum setting. Herein, we present a low-cost, light-based sensor worn on the periphery to monitor for early compensatory mechanisms triggered by blood loss, which was validated in a swine model of hemorrhage and resuscitation. The literature has long described peripheral vasoconstriction as one of the earliest physiologic compensatory mechanisms to protect vital organs from hypo-perfusion and overall cardiovascular collapse^20, 21^. However, this compensation also blunts vital sign responses in the setting of ongoing blood loss, preventing early diagnosis. We demonstrated feasibility of this approach in swine studies, during which we observed significant and near-immediate decreases and increases in LSFI signal during hemorrhage and resuscitation, respectively.

All swine except for the heparinized swine 5 experienced an overall decrease in LSFI during blood loss, indicative of peripheral vasoconstriction in response to hypovolemia. Heparin has been shown to cause vasodilation and lower blood pressure^36, 37^, particularly when administered as a bolus, as it was administered in swine 5. Further, heparin has been shown to reduce endothelial release of the vasoconstrictor endothelin-1^38^ as well as reduce blood viscosity, both of which can affect perfusion. Thus, swine 5 was excluded from quantitative analysis. The quantitative analysis of the correlation between blood loss and LSFI in swine revealed a striking average correlation coefficient of −0.94. Unlike the other non-heparinized swine, swine 2 exhibited an increase in LSFI at the beginning of the hemorrhage procedure followed by a rapid decline. We do expect some biological variability across swine, but also note that swine 2 had the largest amount of IV fluids given prior to the start of the hemorrhage procedure began. It is possible that in this swine, a larger degree of blood loss was required to initiate compensatory response. During resuscitation, all non-heparinized swine showed a net increase in LSFI signal, however swine 3 exhibited a much milder increase than the others. This swine had the greatest volume of blood removed, and the LSFI signal reached a plateau at the last seven blood withdrawals of the hemorrhage phase. It is likely that this swine experienced hypovolemia to the extent of cardiovascular collapse, and the resuscitation fluids provided were not sufficient to re-start cardiovascular flow. A milder response in this case should not be considered a limitation of the technique, rather reflects the true cardiovascular state in response to resuscitation.

In addition to measuring LSFI as a function of net fluid change, we also compared the response of vital signs and SI throughout the hemorrhage and resuscitation procedure. MAP and SI correlated well with blood loss and resuscitation, but LSFI had superior performance in both settings, which supports the utility of using LSFI for hemorrhage detection and management. In these swine studies conducted under general anesthesia, vital signs, particularly MAP, deteriorated quickly with hemorrhage. General anesthesia acts as a vasodilator, and thus compensatory responses where likely blunted. Isoflurane in particular has been shown to inhibit sympathetic response to hypovolemia^39–42^. In future patient studies, we anticipate observing a delay in vital sign deterioration in accordance with the literature^20, 21^, as the vast majority of delivering patients are not under general anesthesia.

It is important to emphasize that during swine studies, an arterial line connected to a SurgiVet Monitor obtained the MAP and recorded data every 6 seconds. In pregnancy, placement of arterial lines is rare due to the high potential for complications; thus, >99% of patients do not have invasive arterial lines. Instead, blood pressure is typically measured using a pneumatic cuff following the oscillometric method, which requires more than 1 minute for a single reading. Our device can currently record at a frequency of 100 Hz, perform frame-by-frame processing at 80 Hz, and transmit data at 1 Hz, allowing for near real-time data visualization and examination of LSFI trends. However we anticipate real-time capabilities (≥30 Hz data processing + transmission) in future iterations. In addition to low sampling rates for blood pressure (BP), studies have demonstrated that oscillometric-based BP cuffs, such as those used in laboring and cesarean delivery patients in hospital settings, perform poorly in hypotensive situations, providing falsely high values that could mask hemorrhage.^43^ It is also crucial to note that LSFI and BP are not measuring the same phenomenon. While the measures are related, they provide distinct information. We observed multiple instances where LSFI and MAP did not correlate (resuscitation phase of swine studies had correlation between LSFI and MAP of 0.36, Supplemental Figure 7), and these situations also demonstrate the benefit of having both pieces of information. For instance, medical decisions would likely differ based on different combinations of MAP and LSFI values. In the case of low MAP and low LSFI (low peripheral perfusion), providers would likely interpret this situation as dangerously low blood volume that needs immediate treatment with either fluids or transfusion. However, in the case of low MAP and normal LSFI, they may instead opt to administer a peripheral vasoconstriction medication to help the patient compensate for mild blood loss and as a result, increase BP, without the need for blood transfusion.

While characterizing our wearable’s performance, we found that, as expected, pairing our low-cost laser with the LSFI wearable device exhibited reduced sensitivity to changes in velocity than the highly coherent benchtop laser and less power stability across a range of ambient temperatures. However, the size and cost of the benchtop laser are prohibitive in being used for wearable applications and the sensitivity to velocity changes was within 75% of the benchmark laser, demonstrating sufficient resolution for the expected in vivo range of velocities. Our wearable system effectively produced a linear response to changes in flow rate, similar to reported findings using high-cost systems,^28^ and successfully detected peripheral perfusion changes in a swine model of hemorrhage and resuscitation, the main criteria for this application. The low-cost laser diode is roughly 12×12×8 mm (width x height x depth) in size, while the benchtop laser, which consists of two separate constant current and temperature drivers that together weigh over 6 kg, require approximately 146×154×320 mm of space.

The Raspberry Pi camera system allowed for speckle-to-pixel ratios that surpassed the Nyquist rate when using 640p at 80 fps and 1920p at 30 fps modes. Swine data was collected while recording at 320p and 100 fps which did not meet the Nyquist sampling criterion, although it approached it. Even without meeting this criterion, the swine data produced promising results, which will improve when the frame rate and consequently pixel binning are reduced. Supplemental Figure 8 demonstrates that decreasing the frame rate will not deleteriously affect the conclusions drawn from the data nor the data quality. With the Raspberry Pi camera, decreasing the frame rate is a simple way to increase speckle size, reduce computation load and needed speed, and improve image quality. A second mechanism to increase the speckle size and the speckle-to-pixel ratio is to decrease the aperture of the camera system, however, this will reduce the amount of light detected by the camera. In addition, reducing the image window size may provide an alternate means to maintain high frame rates (>80 fps) without binning.

Our wearable LSFI sensor is part of a growing group of technologies that measure the physiological response to hemorrhage by interrogating the vasculature. Optical spectroscopy-based sensors tracking longitudinal hemoglobin (Hb) concentration changes caused by postpartum blood loss^44–49^ reported larger bias and decreased accuracy between gold standard Hb measurements compared to similar studies conducted in non-pregnant subjects.^44, 45, 47^ Some authors stated that the devices may need pregnancy-specific calibration prior to successful use in obstetric care. A recent study successfully employed machine learning to improve Hb concentration performance in pregnant patients,^50^ thus, demonstrating that this approach could be further improved for use in pregnancy. One drawback of Hb concentration approaches is that the rate of interstitial fluid transfer to the vasculature is unknown and significant decreases in Hb concentration may take tens of minutes to hours, limiting the utility of this approach in detecting rapid PPH. However, Hb concentration is a principal guide during massive hemorrhage and PPH resuscitation efforts, and therefore is still of clinical value in PPH care.

Researchers have also mined photoplethysmography (PPG) waveforms, the signals acquired from pulse oximeters, to develop predictive algorithms for hemorrhage in military, trauma, and critical care settings.^51^ While these algorithms show promise, PPG has multiple draw backs, including low SNR, particularly in patients with poor perfusion,^52, 53^ as expected during hemorrhage. Furthermore, PPG-based measurements on patients with darkly pigmented skin have lower SNR which can lead to errors when extracting cardiovascular features from PPG waveforms, and could contribute to additional racial disparities in maternal health.^54, 55^

More recently, investigators have begun targeting the peripheral vasoconstriction compensation mechanism as an early indicator of dangerous blood loss. One study evaluated peripheral perfusion using the perfusion index, a pulse oximeter parameter that tracks perfusion based on the ratio of the pulsatile to non-pulsatile signal, to track postpartum blood loss in patients who underwent vaginal delivery. A correlation of −0.7 was found between blood loss and the difference in perfusion index immediately and 20 minutes postpartum.^56^ Although the perfusion index is known to be skewed and has high patient variability,^57^ these results show that non-invasive measures of peripheral vasoconstriction can provide early signs of postpartum blood loss. An alternate strategy to the perfusion index is using dynamic light scattering (DLS) to monitor blood flow. A miniaturized DLS (mDLS) system was developed^58^ and tested in a swine hemorrhage model, with

Our wearable LSFI sensor has numerous advantages over the aforementioned methods. LSFI has demonstrated significantly higher SNR than PPG^29, 59^ which measures changes caused by light scattering rather than PPG measurements of absorption. Furthermore, tests quantifying the effects of skin pigmentation in both phantom measurements and patient studies found no significant difference based on level of pigmentation.^31^ LSFI has a broad dynamic range with linear response to blood flow even in low perfusion conditions^29^), which overcomes limitations of the PPG-derived from the perfusion index. While LSFI and mDLS share similarities in what is being measured, by using a camera consisting of thousands of photodetectors instead of a single photodetector, LSFI provides a higher SNR with relatively simple and low-cost instrumentation.

Current limitations of the device include the fact that the device measures relative values of blood velocity/flow rather than absolute measurements. This is a limitation that affects all laser speckle-based imaging devices and is an active area of research within the laser speckle imaging community. Some advancements have been made on this front, in which using a wide range of exposure times allows for a more accurate estimation of blood velocity.^60^ In addition, the current device is relatively bulky and has high power consumption, which will both need to be reduced in future iterations. Additional features should allow for optimization of the device parameters for each experiment, helping to ensure high SNR and large velocity dynamic range. Such features include autofocus and adjustable light intensity via altering exposure time or varying laser power until a desired light level is reached and then set for the duration of the experiment. The largest limitation of the current device is laser instability, which can be corrected by developing more sophisticated laser driver circuitry and testing new compact and inexpensive laser diode options.

In summary, we have demonstrated that a custom wearable laser speckle imaging device could rapidly detect hemorrhage-driven peripheral vasoconstriction and resuscitation via crystalloid infusion. This progress provides a framework for a novel low-cost and noninvasive technology that identifies ongoing blood loss when low-cost and accessible interventions are effective, with the hope of reducing the unacceptably high rates of maternal morbidity and mortality caused by hemorrhage in low- and high-resource settings alike.

## Methods

### 1. Sensor hardware design

The wearable laser speckle-based peripheral perfusion sensor was configured in reflectance mode, with the excitation source consisting of a 50 mW 785 nm laser diode (Laserland, 11071013). This wavelength was chosen because near infrared sources have higher tissue penetration depth and minimum melanin absorption (Zonios et al., 2008), thus reducing skin pigmentation-based signal bias while still operating at a wavelength that is sensitively detected by inexpensive silicon-based CMOS detectors. The center-to-center distance between the CMOS sensor and laser was set to 11 mm, enabling collection of diffusely scattered photons that have interacted with subsurface blood vessels^30^. The sensor system consists of a double-lens Raspberry Pi camera Module v2 NoIR. This camera has an 8MP Sony IMX219 CMOS sensor and no infrared (IR) filter, allowing the camera to detect IR light. The camera is capable of acquiring 3280×2464 pixel images and recording video at 1080p@30 frames per second (fps). For this study, recording at 320×240p allowed 100 fps, and 80 fps was possible at 640×480p. The sensor size is 3.68 × 2.76 mm, with each pixel being 1.12 × 1.12 µm.

A double-lens system was constructed by mounting the original Raspberry Pi camera fixed focus lens^61^ on the module and introducing a second identical lens in an inverted position placed at the surface of the first lens. A 0.4 mm thick, 7.5 mm diameter sapphire optical window (Edmond Optics, 43-628) was placed on the surface of the second lens, allowing the system to be in focus when an object is in direct contact with the window. This configuration enabled the microscopic resolution from a short optical train^62^ of 6.42 mm. Such miniaturization is ideal for wearable devices which require a small footprint.

The device is controlled by a Raspberry Pi Zero 2W computer and powered by either wall power or a PiSugar 1200 mAh battery (Guangzhou Soft Electric Technology Co., Ltd., Guangzhou, China). The Pi Sugar battery is an uninterruptible power source that can supply power and charge simultaneously, allowing the device to be wireless while maintaining voltage regulation. A custom PCB provided a stable 3.3 V to the laser diode from a constant 5 V pin on the Raspberry Pi computer and a general-purpose input/output (GPIO) pin was used for laser control.

### 2. Sensor software design

The device software, written in Python, controlled the laser diode and camera functionality. The Raspberry Pi computer interfaced with a PC laptop via Bluetooth connection (Figure 1). In this connection, the Raspberry Pi was the server and the PC was the client. The PC took user input to turn on/off the laser diode and record laser speckle videos for a specified length of time. The Raspberry Pi carried out desired tasks, saved recorded speckle videos, processed video data, and sent processed data to the PC. The PC received, saved, and plotted data as a function of time. This program allowed for near real-time data visualization for monitoring blood flow.

To extract LSFI, laser speckle contrast (K) was first computed on the Raspberry Pi Zero 2W computer using a custom algorithm for fast processing. Each frame from the video was processed using the spatial processing algorithm to calculate the laser speckle contrast image.^63^ A 7×7 sliding window translated across a raw intensity image to produce a speckle contrast value (K) for each window within the frame:

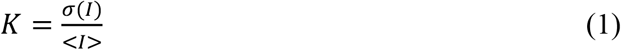

where *σ*(*I*) is the standard deviation of pixel intensity within the window and < *I* > is the mean pixel intensity within the window. Following the mathematical process outlined by Tom et. al.,^64^ equation 1 can be rewritten in terms of summations of pixel intensities and squared pixel intensities:

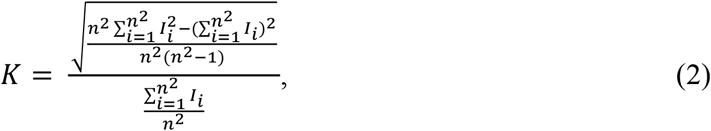

where *n x n* is the dimensions of the sliding window. This is equivalent to convolving the raw image and square of the raw image with an *n*x*n* matrix of ones. We found that computation speed can increase while retaining the speckle contrast value by convolving with an *n*x*n* identity matrix:

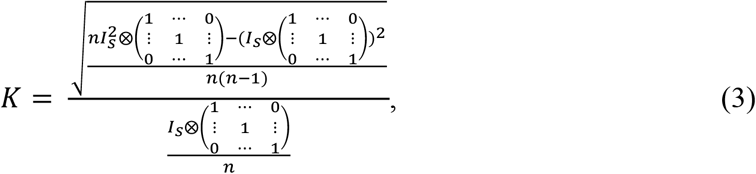

where *I*_*s*_ is the raw image and 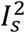 is the square of the raw image. An efficient computational approach to calculate these convolutions is to generate ‘diagonal speckle sums’ *I*_*ds*_ and 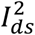 for both the raw image and the square of the image respectively:

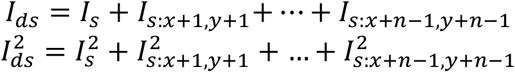

where *I*_*s:x*+*k,y*+*k*_ and 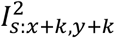 are the raw image and square of the raw image offset *k* pixels along the diagonal. The diagonal speckle sums can then be used instead of the convolutions to generate a pixel-by-pixel speckle contrast value via the following equation:

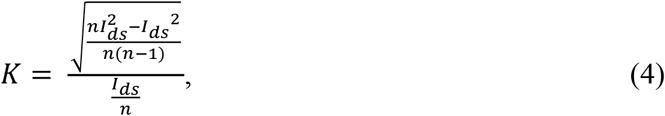

The mean value of *K* across all imaging pixels defines ⟨*K*⟩. The Raspberry Pi computer implemented the algorithm in python and transmitted ⟨*K*⟩ to the receiving PC. ⟨*K*⟩ was then converted to the laser speckle flow index (LSFI), which is a measure proportionate to blood flow (perfusion), via the following equation:

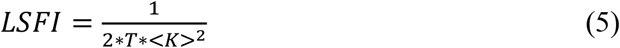

where *T* is the camera exposure time. This process was repeated for every frame of the laser speckle video, resulting in a time-varying pulsatile waveform known as the speckle-plethysmogram (SPG).^29^ A wide moving average filter applied to the waveform removed pulsatile variation and produced a clean mean LSFI signal.

### 3. Impact of camera system parameters on laser speckle

High-contrast laser speckle imaging requires proper spatial or temporal sampling. Spatial sampling provides higher temporal resolution to observe rapidly changing phenomena such as blood flow. Optimization of the imaging system allows for observation of the full laser speckle pattern. The minimum speckle size should be greater than two times the pixel size to surpass the Nyquist sampling rate. Speckle size can be calculated using the equation:

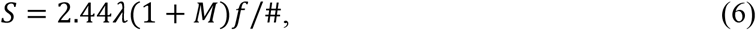

in which *λ* is the wavelength of the coherent light source, *M* is the magnification of the camera system, and *f*/# is f number of the camera system.^63^

Capturing 1920×1080, 640×480, and 320×240 pixel videos at 30, 80, and 100 fps, respectively, of a resolution test target (Thorlabs, R1DS1P) enabled determination of the double-lens camera system magnification by comparing the physical pixel size to the object size per pixel:

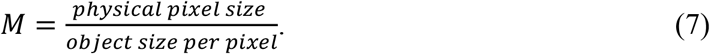

The object size per pixel was calculated using MATLAB. Physical pixel size is a characteristic of the sensor, however, when the resolution is reduced to 640×480 or 320×240 under default settings, the camera sensor performs 2×2 or 4×4 pixel binning, increasing the pixel size by a factor of 2 or 4, respectively.

The f/# of the system was found in the Raspberry Pi v2 camera module documentation, and experimentally verified using the equation:

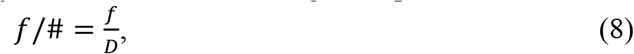

where *f* represents the focal length of the system and *D* represents the diameter of the lens. The Raspberry Pi documentation provided the *f*/# and *f* of the single lens system which were used to solve Eq. 8 for the diameter of the lens which was verified by physical measurement of the lens aperture.

The spatial resolution of the system was found by plotting the pixel intensities of a row that spanned across a set of 3 lines on an USAF resolution test target (Thorlabs) and ensuring a step function was observed. This process was performed on multiple sets of lines until the smallest line thickness that produced a step function was determined.

### 4. Laser characterization

To ensure laser stability over time while in use, the spectral profile and power of the low-cost laser were recorded every 10 minutes over a six-hour period using a spectrometer (Flame Spectrometer, Ocean Optics) and an optical power meter (PM100D and S121C, Thorlabs). An identical experiment conducted for benchmarking performance used a high-cost laser system that included a pigtail laser (LP785-SAV50, Thorlabs), laser mount (LDM9LP), temperature controller (TED 200, Thorlabs), and current driver (LDC 205 C, Thorlabs), and cost ∼100X more than the low-cost laser diode. In addition, recording the spectral profile and power of the low-cost laser at temperatures ranging from 13-40*°*C, representing the range of possible ambient temperatures a patient might experience, provided the stability of the laser with varying temperatures. Operating the lasers within a 0*°*C cold room lower the temperature, while warm air from a heat gun (840015, Kawasaki) raised the temperature. An infrared camera (E6-XT, FLIR) monitored the temperature. Measuring the mean LSFI over a 15 second period while recording a static piece of white paper (minimum) or a blood flow phantom at a 20 mm/s (maximum) produced the distribution of pixel intensities and the range of speckle values for each laser. The LSFI was measured at 100 fps with an exposure time of 5 ms to mirror the parameters used in the swine hemorrhage model.

### 5. Device power consumption

The power consumption of each device during various states were measured using a digital multimeter (Eversame, 00005). Power consumption was monitored during idle states, data collection, and data transmission modes. Additionally, the electric charge (ampere-hours) consumed by the device over a 30-minute recording period was measured to determine the battery capacity required to power the device.

### 6. Device performance in vitro

In vitro testing was performed using an optical tissue-mimicking flow phantom. Custom optical flow phantoms were created using an Elegoo Saturn S MSLA 9.1” 4K mono LCD resin-based 3D printer and translucent ABS-Like Photopolymer resin (Elegoo, Shenzhen, China). The optical phantom was designed as a CAD model in Fusion 360 (Autodesk, San Rafael, CA) to be a rectangular solid with 20 mm length and 10 mm width and height. A 1.6 mm diameter horizontal channel was modeled lengthwise through the phantom, centered and 2 mm from the top face. The CAD model underwent slicing via ChituBox V1.9.3 for printing. To ensure that the channel did not collapse during the print, the 3D print was paused in the middle of the print process, at which time the phantom consisted of a rectangular solid with a cylindrical groove at the top of the print. Then a 1.45 mm diameter (15 American Wire Gauge, or AWG) wire was placed along the groove and secured tightly around the build plate. The wire used was slightly smaller than 1.6 mm to allow tolerance for channel shrinkage (allowing the channel to form around the wire) and to protect the LCD screen of the printer from coming into direct contact with the wire. The wire insertion process occured quickly to reduce the risk of layer separation due to the resin cooling, and a heat gun warmed the resin before resuming the print with the wire in place. After printing, the wire was cut from the build plate and pulled out gently from the print. The model was then rinsed in ethanol and post-cured in a UV station for 1 minute facing upward and 1 minute facing downward.

A syringe containing 2% intralipid fluid (l141, Sigma-Aldrich, St. Louis, MO) was connected to the hollow channel of the optical flow phantom using a luer lock needle. A syringe pump (Ne-1000, New Era Pump Systems Inc.) pushed fluid through the optical phantom at various physiologically relevant velocities ranging between 0.75 and 22.5 mm/s. Fluid was pumped for 20 seconds at each velocity and laser speckle video was recorded for the last 3 seconds of each interval to allow the channel pressure to equilibrate at each velocity. Images were acquired at an exposure of 0.5 ms and 80 fps, resulting in laser speckle images that surpassed Nyquist criteria for speckle size. The mean LSFI over the 3 second interval was calculated for each velocity.

### 7. Device performance in swine model of hemorrhage

All swine studies were performed under an approved IACUC protocol with supervision by veterinary staff at Washington University in St. Louis, Division of Comparative Medicine. The study used 11-week-old male Yorkshire and Landrace cross-breed pigs (Oak Hill Genetics, Ewing, IL). The average weight of the pigs was 37.09 kg and the range of weights was 33.57-43.18 kg.

All pigs had a standard laboratory diet with ad-lib access to water prior to beginning the experiments. Pigs were fasted from food for 12 hours prior to the start of the procedure. All hemorrhage studies were acute, with all pigs euthanized using intravenous (IV) potassium chloride immediately after the completion of hemorrhage and volume resuscitation procedures.

A telazol/ketamine/xylazine cocktail (TKX) given intramuscularly sedated the swine, followed by intubation with a 6-or7-mm endotracheal tube. Isoflurane gas maintained the surgical plane of anesthesia for the animals and prevented pain, distress, or movement during the experiments. To monitor vital signs throughout the studies, a thermometer tracked temperature, and a Surgivet® Advisor® Tech Vital Signs Monitor (Smiths Medical, Plymouth, MN) tracked heart rate and SpO_2_ via a pulse oximeter applied to the swine tongue and arterial blood pressure via invasive pressure catheters. An ear vein catheter provided the animals with IV fluids and intraoperative medications. All animals received IV fluids at a drip rate of 3-6 ml/min during the procedure. Prior to initiation of the study, the animals were placed in a supine position on a heated operating table and maintained on positive pressure ventilation (10 ml/kg, 12 breaths per minute). Blood glucose was maintained >60 mg/dL via supplemental dextrose IV as needed. Swine 4 experienced several blood clots in the femoral catheter line, making blood collection difficult. The next pig (swine 5) was heparinized (150 IU/kg IV) to prevent blood clotting; however, effects on peripheral perfusion were noted (hypotension and non-reactive peripheral vessels in response to hemorrhage). Subsequent pigs were not heparinized. Instead, a heparinized saline (6 IU/ml) lock was maintained in the large bore pressure catheters.

In the first swine, placement of vascular access sheaths (7F) in the left femoral vein and right femoral artery was performed using surgical dissection. Subsequent vascular access sheaths were placed in the other pigs using either surgical dissection or an ultrasound-guided percutaneous technique. The left femoral vein sheath was used for blood removal to induce hemorrhagic shock, however, the vein collapsed during blood removal. In all subsequent swine, blood was withdrawn from the left femoral artery, as the artery is less likely to collapse during blood removal due to more muscular tone. The right femoral catheter measured systolic, diastolic, and mean arterial pressure (MAP) via the Surgivet® Adivsor® Monitor. Three study pigs also had a vascular access sheath (7F) placed in the right external jugular vein via surgical dissection. Catheters placed in the external jugular vein measured central venous pressure (CVP) and patency was maintained via heparinized saline as described above. Experienced laboratory animal veterinarians inserted all catheters.

Preparation for the placement of the laser speckle device included shaving the left posterior carpus of the swine, which minimized interference with the sensor and affixing the sensor to the shaved area after sedation. A minimum 20-minute baseline recording occurred before initiating the hemorrhage protocol. The LSFI recorded at 100 fps for 10 seconds of each minute and calculated the mean over each 10-second interval. A 5-frame moving average helped to smooth the data collected during the study. The LSFI from each swine was normalized to the highest LSFI value obtained throughout the study (peak-normalized) and plotted against net fluid change.

The controlled volume hemorrhage protocol was followed by removing either 1.5 or 3% EBV every 2.5 or 5 minutes until either a total 40% EBV had been removed or MAP neared 40, either of which defined reaching hemorrhagic shock. Once animals experienced hemorrhagic shock, blood withdrawal ceased and no intervention were given for 10 minutes. After the 10-minute pause, IV infusion with crystalloids (NaCl saline solution or Lactated Ringers Solution) at 1.5-3% EBV occurred every 5 minutes for a total of 30 minutes. EBV was calculated based on the approximation that each pig has an EBV_min_ of 58 mL blood / kg and an EBV_max_ of 74 mL blood / kg.^65^ Throughout the procedure, blood was collected in heparinized 1 mL syringes and used to measure blood gas, hematocrit, sodium, chlorine, calcium, glucose, and lactose contents of the blood in 15 minutes intervals. Vital signs including heart rate, core temperature, respiration rate, CVP, and blood oxygen saturation were also monitored by veterinary staff and included in the study results.

### 8. Laser speckle video parameter analysis

Beyond the pixel size requirements discussed above, numerous additional camera parameters can affect the quality of laser speckle contrast, including camera frame rate, exposure time, and frame size (number of pixels). The frame rate used to record video is the sampling frequency of the system. It is essential to sample at a rate that allows for the signal to be reconstructed without loss of vital blood flow information. As is typical for most imaging systems, there is a tradeoff between the frame rate and the image quality. At higher frame rates, the Raspberry Pi camera will reduce to a smaller field of view, utilizing fewer pixels on the sensor, and will default to pixel binning. Comparing a video recorded at 100 fps with the same down-sampled video at 50 and 10 fps enabled examination of the effects of frame rate on LSFI information content. Furthermore, performing the LSFI calculation on each full, half, and quarter frame allowed the impact of frame size to be studied. The half and quarter frames had the same aspect ratio as the full frame and utilized the centermost pixels. This method permitted determination of the minimum necessary sampling rate and image size.

## Acknowledgments

The authors gratefully acknowledge funding from the National Institute of Child Health and Human Development (NICHD, K99HD103954 & R00HD103954), the Washington University in St. Louis Collaboration Initiation Grant, and helpful discussions and feedback from Joe Culver, Sam Achilefu, Sarah England, David Monks, Patricia Strutz, Mike Dombrowski, Roxane Rampersad, Deborah Frank, and Molly Stout.

