## Supplemental Figures for "Development and evaluation of a wearable peripheral vascular compensation sensor in a swine model of hemorrhage"

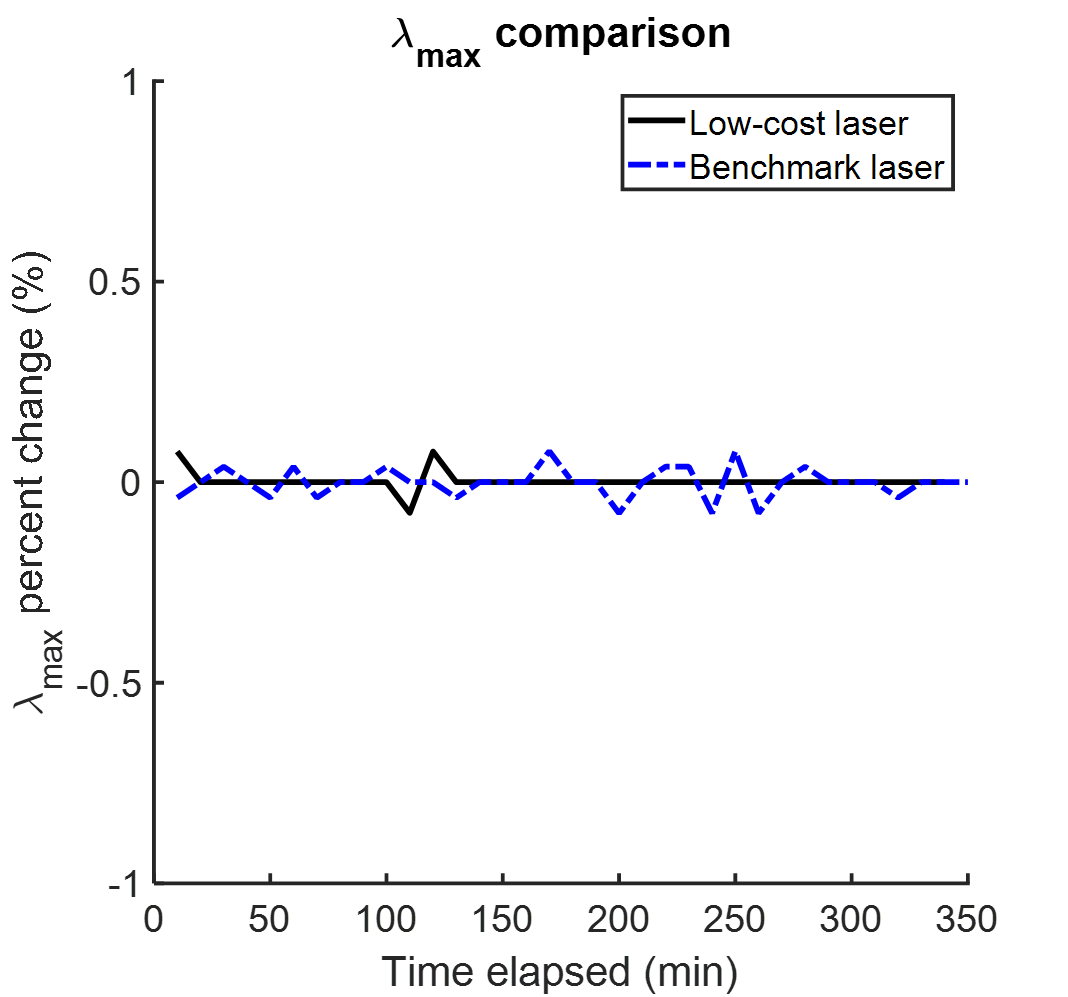

***Supplemental Figure 1.*** *Percent change of peak wavelength in low-cost and benchmark laser.*

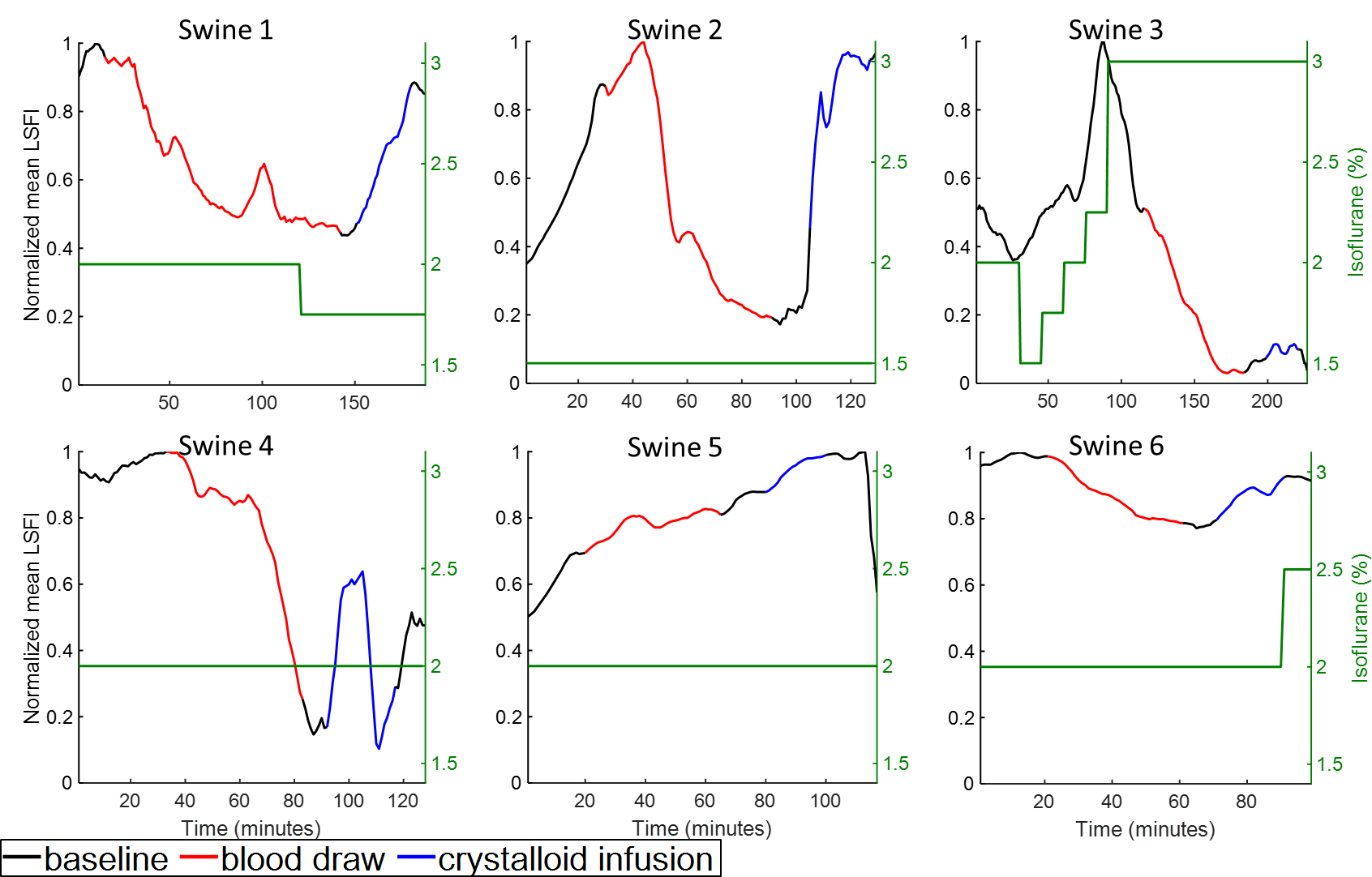

***Supplemental Figure 2.*** *Peak-normalized LSFI signal with isoflurane dose overlaid.*

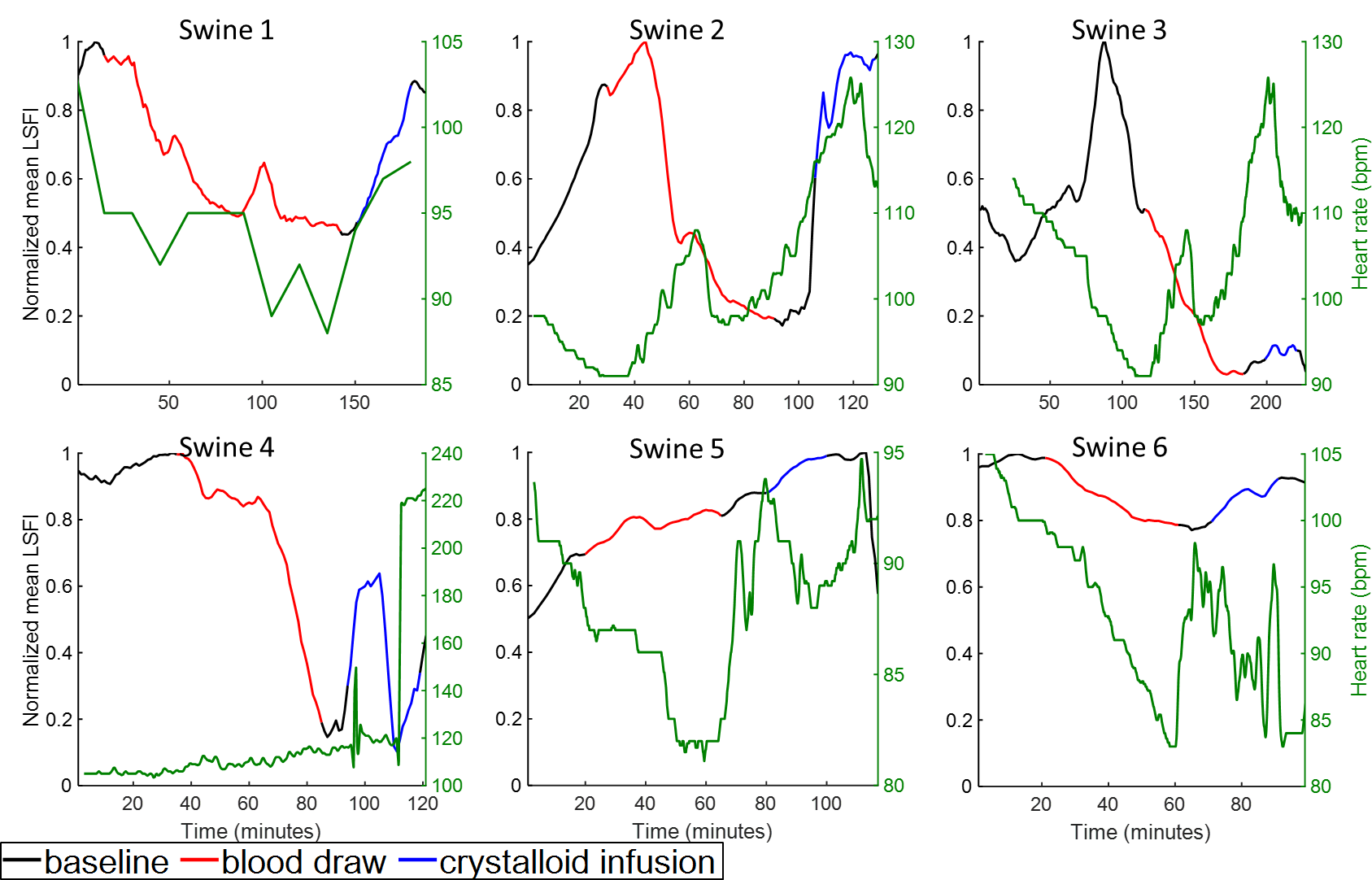

***Supplemental Figure 3.*** *Peak-normalized LSFI signal with heart rate overlaid.*

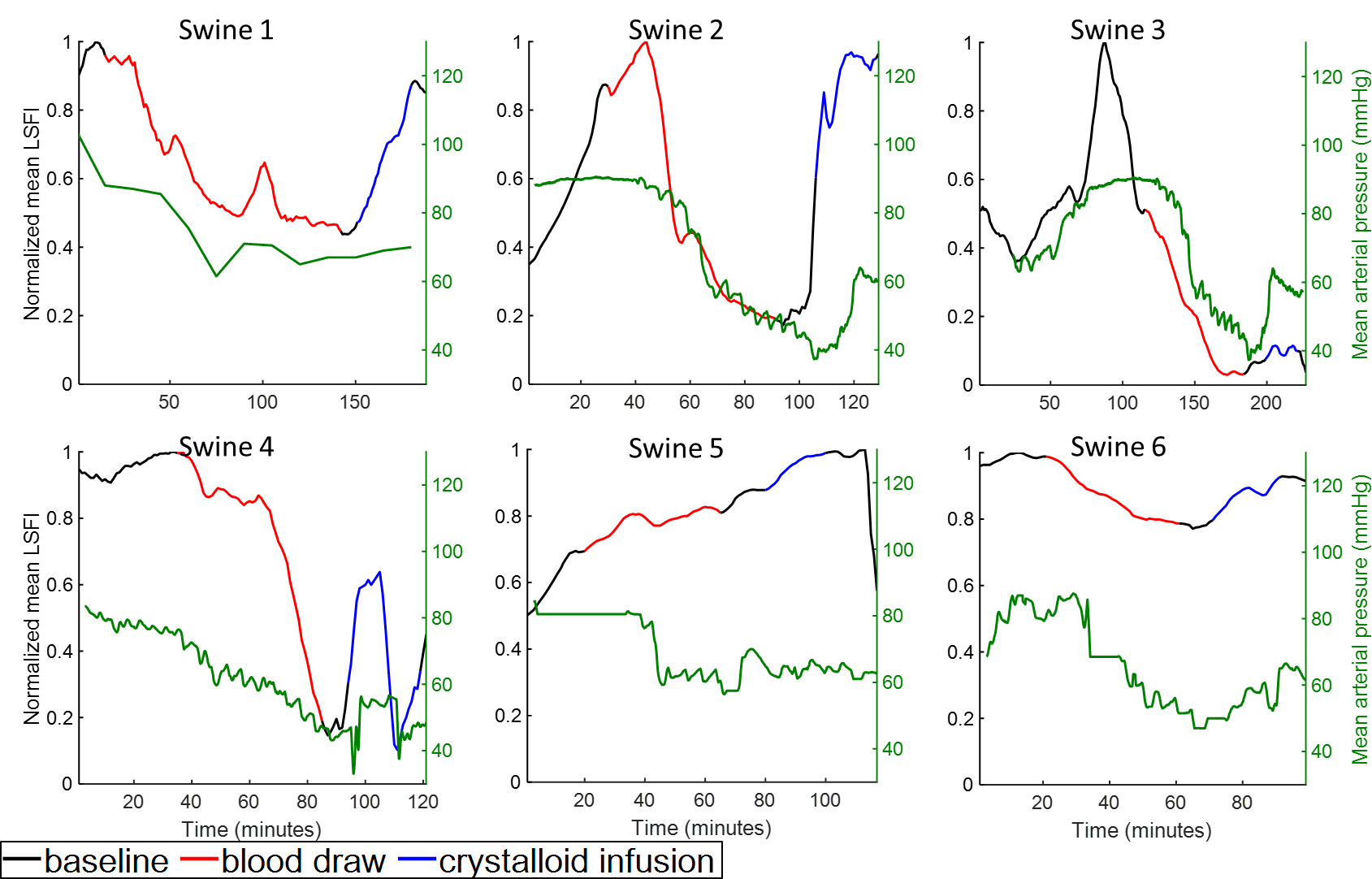

***Supplemental Figure 4.*** *Peak-normalized LSFI signal with mean arterial pressure overlaid.*

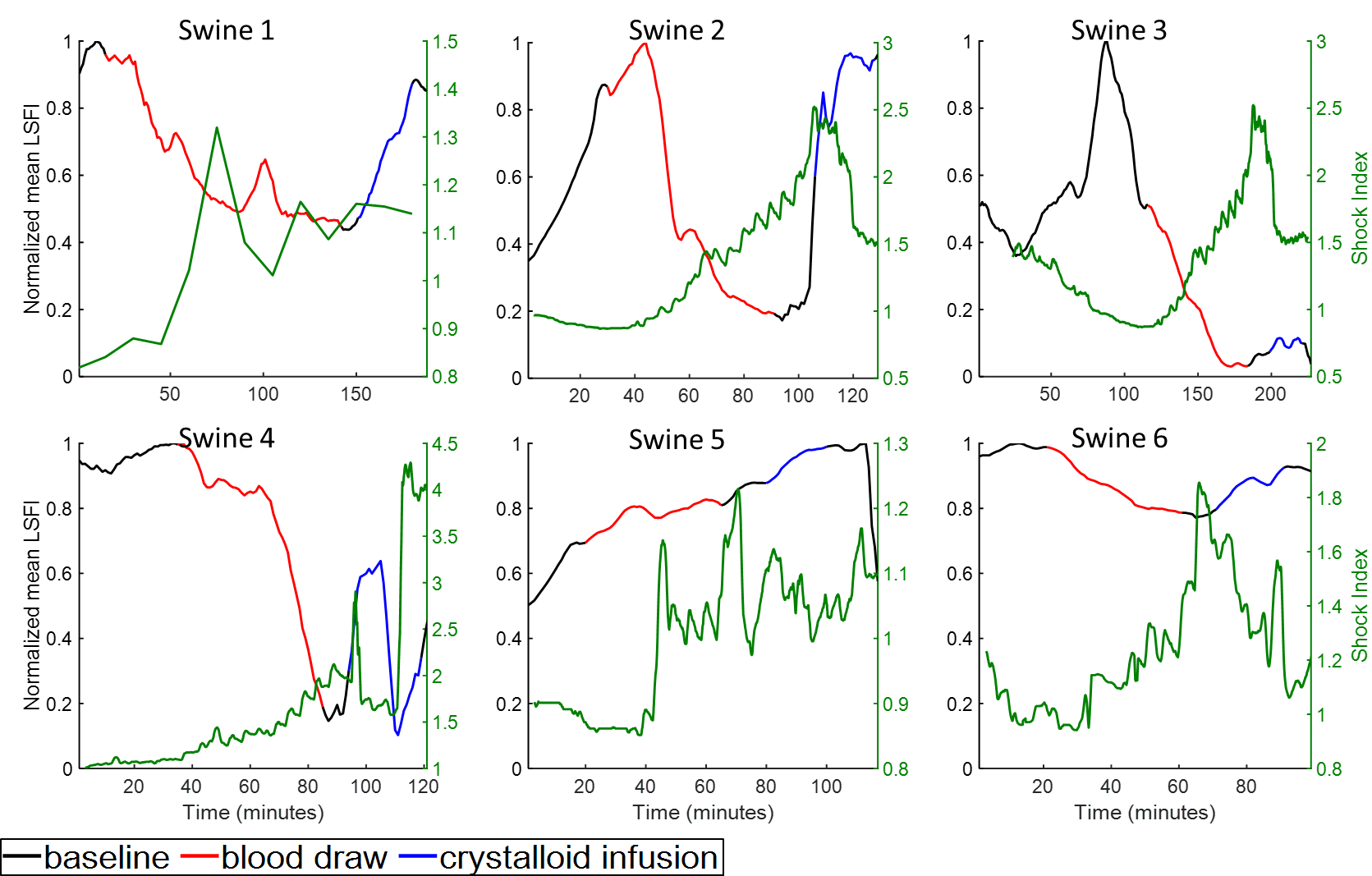

***Supplemental Figure 5.*** *Peak-normalized LSFI signal with Shock Index overlaid.*

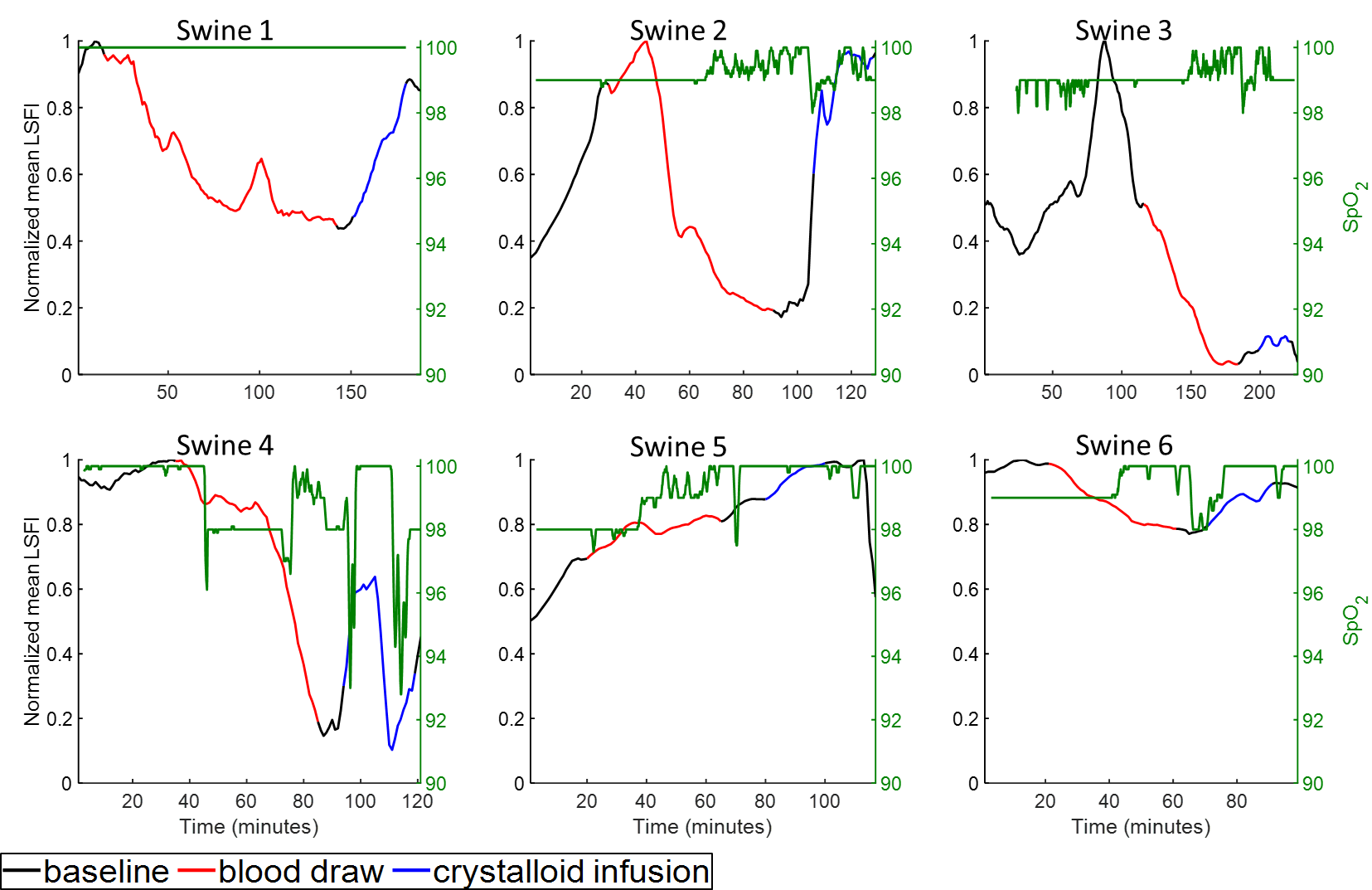

***Supplemental Figure 6.*** *Peak-normalized LSFI signal with blood oxygen saturation (SpO_2_) overlaid.*

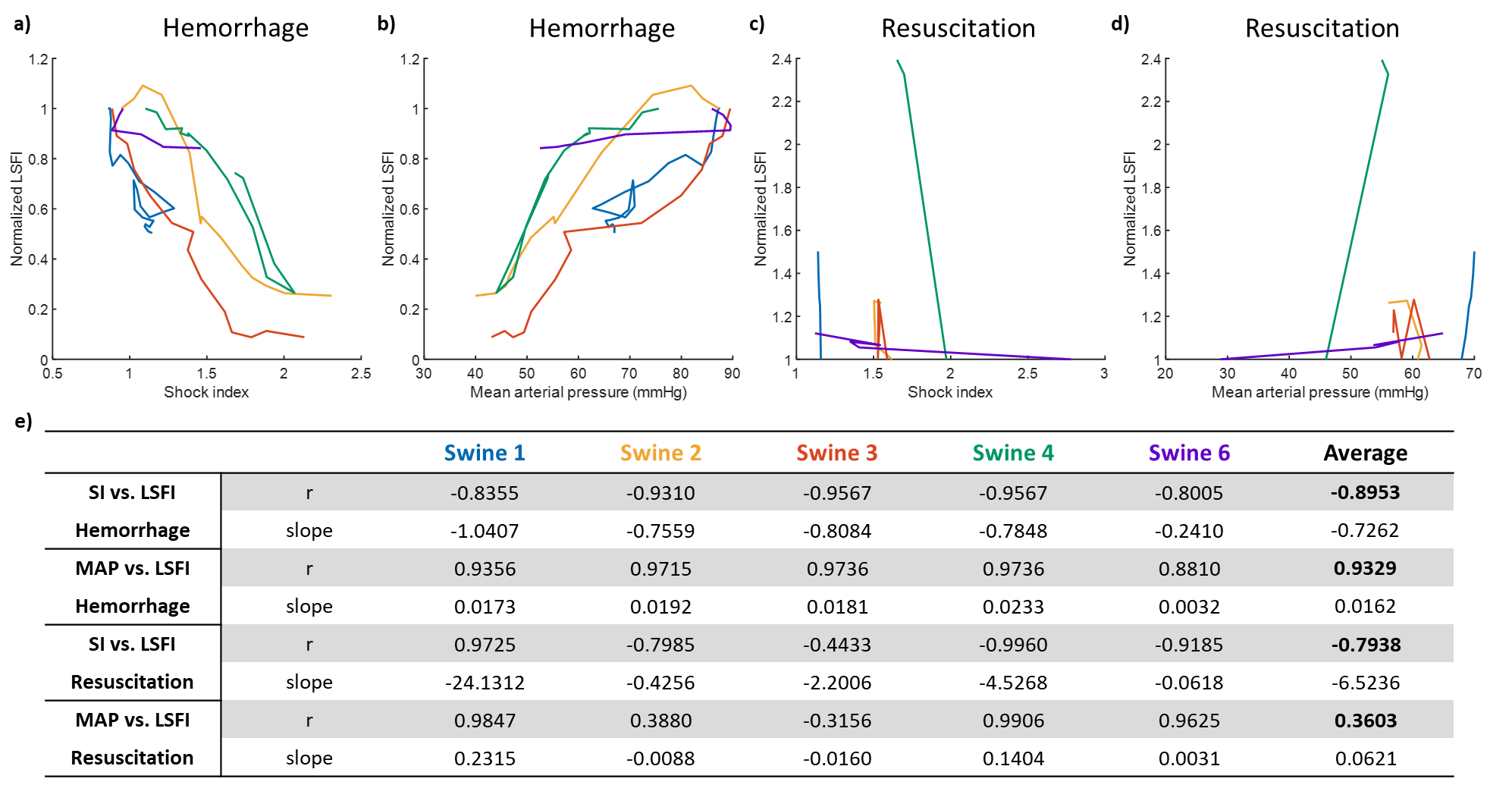

***Supplemental Figure 7.*** *Comparison of SI vs. normalized LSFI and MAP vs. normalized LSFI during hemorrhage (a-b) and resuscitation (c-d). Summary of correlation coefficients and slopes for each linear correlation.*

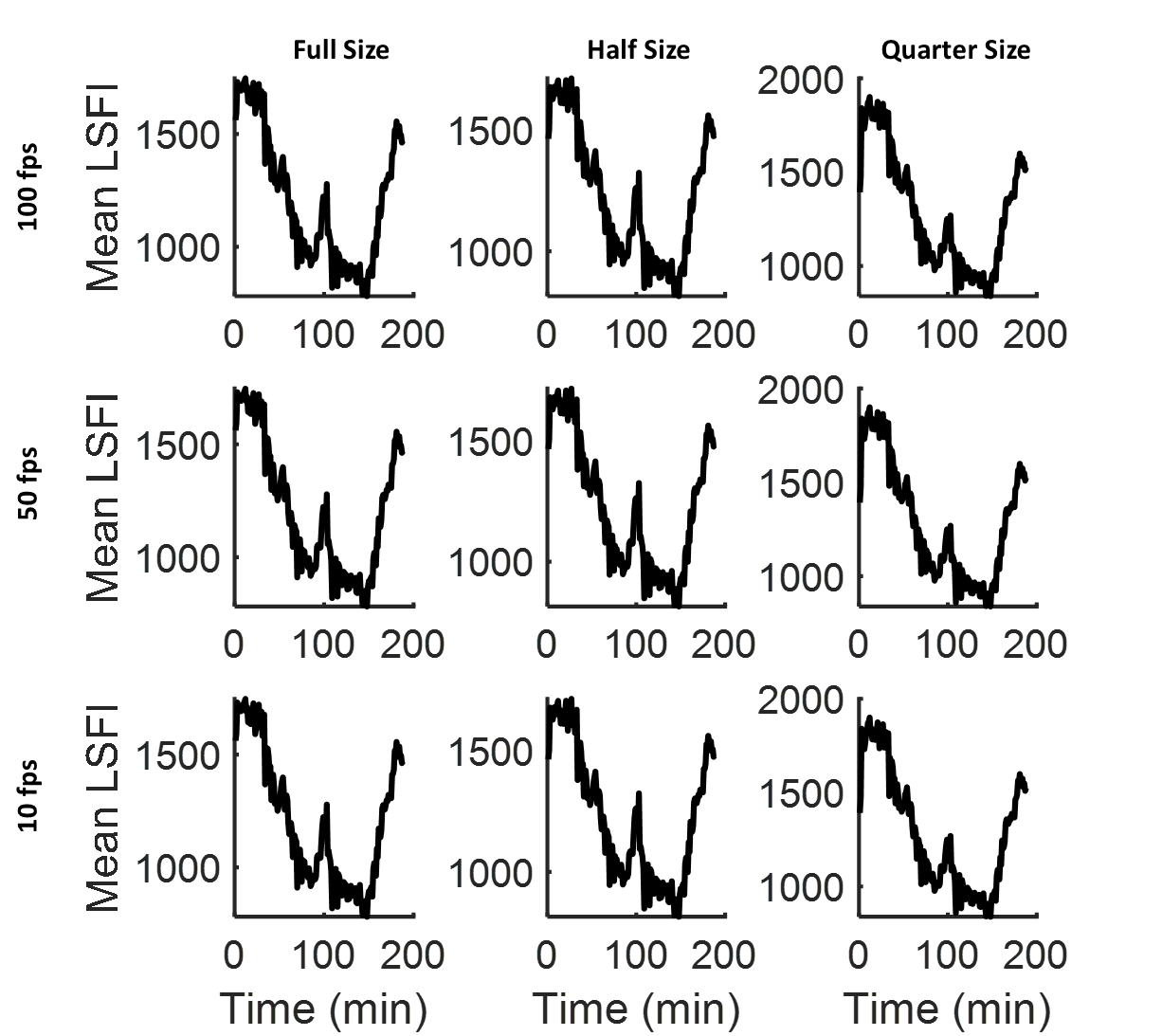

***Supplemental Figure 8.*** *Effects of sampling rate and frame size on mean LSFI signal in swine 1.*
